# Strength of T cell signaling regulates HIV-1 replication and establishment of latency

**DOI:** 10.1101/432401

**Authors:** M Gagne, D Michaels, GM Schiralli Lester, WW Wong, S Gummuluru, AJ Henderson

**Author notes:** Corresponding author, (AJH).

## Abstract

A major barrier to curing HIV is the long-lived latent reservoir that supports re-emergence of HIV upon treatment interruption. Targeting this reservoir will require mechanistic insights into the establishment and maintenance of HIV latency. Whether T cell signaling at the time of HIV-1 infection influences productive replication or latency is not fully understood. We used a panel of chimeric antigen receptors (CARs) with different ligand binding affinities to induce a range of signaling strengths to model differential T cell receptor signaling at the time of HIV-1 infection. Stimulation of T cell lines or primary CD4+ T cells expressing chimeric antigen receptors supported HIV-1 infection regardless of affinity for ligand; however, only signaling by the highest affinity receptor facilitated HIV-1 expression. Activation of chimeric antigen receptors that had intermediate and low binding affinities did not support provirus transcription, suggesting that a minimal signal is required for optimal HIV-1 expression. In addition, strong signaling at the time of infection produced a latent population that was readily inducible, whereas latent cells generated in response to weaker signals were not easily reversed. Chromatin immunoprecipitation showed HIV-1 transcription was limited by transcriptional elongation and that robust signaling decreased the presence of negative elongation factor, a pausing factor, by more than 80%. These studies demonstrate that T cell signaling influences HIV-1 infection and the establishment of different subsets of latently infected cells, which may have implications for targeting the HIV reservoir.

**Author Summary:** Activation of CD4+ T cells facilitates HIV-1 infection; however, whether there are minimal signals required for the establishment of infection, replication, and latency has not been explored. To determine how T cell signaling influences HIV-1 infection and the generation of latently infected cells, we used chimeric antigen receptors to create a tunable model. Stronger signals result in robust HIV-1 expression and an inducible latent population. Minimal signals predispose cells towards latent infections that are refractory to reversal. We discovered that repression of HIV-1 transcription immediately after infection is due to RNA polymerase II pausing and inefficient transcription elongation. These studies demonstrate that signaling events influence the course of HIV-1 infection and have implications for cure strategies. They also provide a mechanistic explanation for why a significant portion of the HIV latent reservoir is not responsive to latency reversing agents which function by modifiying chromatin.

## Introduction

HIV-1 persists in a transcriptionally silent latent state in long-lived memory T cells. Although antiretroviral therapies (ART) suppress HIV-1 replication, interruption of treatment results in rapid viral rebound. Therefore, HIV-1 patients must remain on ART indefinitely, despite long term side effects, development of treatment resistance, and viral-induced inflammation [1–3]. For this reason, one strategy currently being explored for cure efforts is “shock and kill,” in which latent HIV-1 is reactivated in conjunction with ART using latency-reversing agents (LRAs). Following reactivation, infected cells are predicted to be eliminated by HIV-specific immunity or virally induced apoptosis. However, clinical trials using LRAs have only minimally perturbed the size of the viral reservoir [4–6].

A cure for latent HIV-1 will require a better understanding of the biochemical factors involved. Latency in chronically infected primary cells and cell lines is regulated by multiple transcriptional mechanisms including NF-κB activation, chromatin accessibility, provirus transcription initiation, Tat availability, P-TEFb sequestration, and transcriptional elongation [7–11]. However, what is not understood is how latency is initially established within a cell and if events at the time of HIV-1 infection influence the transcriptional status of the provirus. These questions are relevant since the latent reservoir is established within the first two weeks of infection [12,13]. New cure strategies will need to limit the size of the reservoir at early time points.

One mechanism that could predispose HIV-1 towards active replication or transcriptional repression and latency is signaling through the T cell receptor (TCR). Engagement of the TCR and costimulatory CD28 molecule result in a multitude of cellular outcomes that influence HIV replication including cytoskeleton reorganization, the activation of transcription factors, enhanced RNA polymerase II (RNAP II) processivity, and chromatin remodeling [14–16]. We hypothesized that the magnitude of T cell signaling during HIV-1 infection will dictate the course of the infection. In order to manipulate signal strength received by a T cell at the time of HIV-1 infection, we utilized chimeric antigen receptors (CARs) that recapitulate T cell receptor and CD28 signaling. By modulating the affinity with which these CARs bind to their ligand, we can differentially deliver signals to target cells.

Using these CARs, we demonstrate that stronger T cell signaling at the time of HIV-1 infection increases subsequent HIV-1 transcription and replication. Robust signals also facilitated the formation of latently infected cells that were readily inducible upon secondary stimulation. Minimal signaling through CARs, although sufficient for HIV-1 integration, failed to support viral replication and generated a deep-seated latent infection. Transcriptional elongation of HIV-1 provirus was limited by RNAPII pausing in the absence of CAR signaling; however, strong CAR signaling correlated with decreased negative elongation factor (NELF) binding and enhanced RNAPII processivity. Our results suggest a model in which signaling strength influences HIV-1 transcription and establishment of latency at the time of initial infection of CD4+ T cells.

## Results

### CARs induce T cell signaling

To examine how signaling cascades downstream from the T cell receptor regulate HIV-1 transcription we utilized CARs (Fig 1A). Intracellular signaling domains for the CARs include CD3ζ with its immunoreceptor tyrosine-based activation motifs (ITAMs) and the CD28 costimulatory domain with its four critical tyrosine residues [17]. Furthermore, a mCherry tag provides a marker for positive selection of CAR+ cells. The extracellular ligand-binding domains of the CARs consist of a single chain variable fragment (scFv) that recognizes receptor tyrosine-protein kinase erbB-2 (Her2) [18,19]. By using different scFvs, a library of CARs with binding affinities for Her2 ligand spanning three logs were generated (Fig 1B). CARs were transduced into Jurkat T cells and primary CD4+ T cells. By enriching for mCherry, we obtained CAR+ populations that were > 90% pure (Fig 1C).

**Fig 1.**
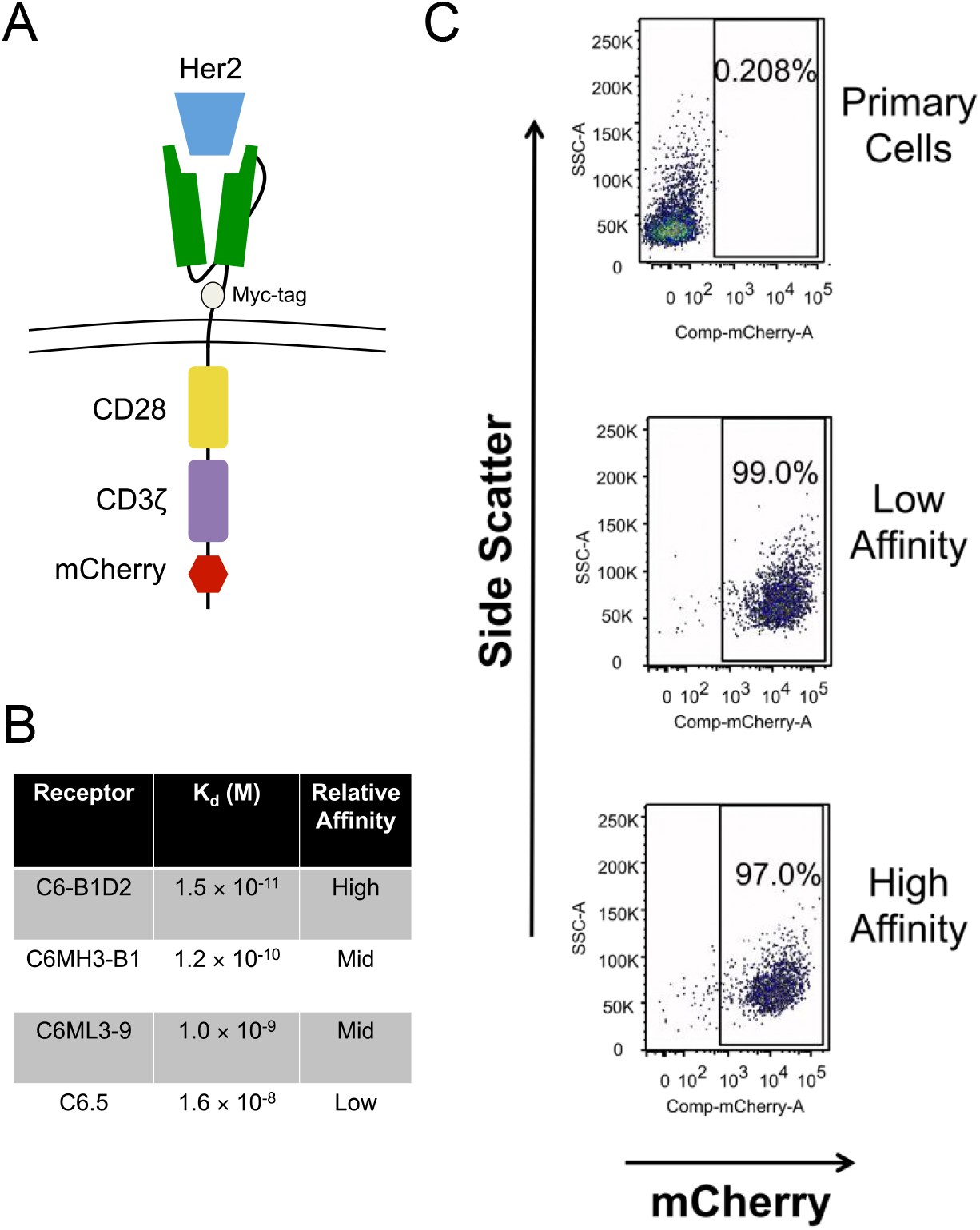
Chimeric antigen receptors used for tunable T cell signaling. *(A)* The design of CARs including CD3ζ and CD28 signaling domains. (*B*) CAR single chain variable fragments with their corresponding dissociation constants for the Her2 ligand. We refer to these CARs by their relative ligand affinity. (*C*) Enrichment of primary CD4+ T cells based on mCherry expression following CAR transduction. See also S1 Fig.

To confirm that signaling through the CARs mimicked TCR signaling, CD69, a transmembrane lectin and a marker for CD4+ T cell activation, was monitored by flow cytometry before and after receptor activation with Her 2 ligand (S1 Fig). Primary CD4+ T cells transduced with either the low affinity or the high affinity receptors were stimulated with plate-bound Her2 ligand for 24 h. In the absence of ligand, less than 7% of the cells were positive for CD69, verifying that there is no ectopic CAR signaling. Activating cells with Her2 induced CD69 expression in the low affinity and high affinity receptors relative to their affinity for ligand. These data demonstrate that CARs can be used as a tool to modulate T cell signaling.

### T cell signaling at the time of HIV-1 infection regulates provirus expression

To determine whether T cell signaling influences viral infection, Jurkat T cells expressing low affinity or high affinity CARs were plated on Her2-coated wells and simultaneously infected with VSV-G pseudotyped NL4–3.Luc, a single-cycle HIV-1 clone which contains a luciferase reporter in place of Nef. VSV-G allowed us to bypass potentially confounding effects from receptor/chemokine receptor signaling due to gp120 binding and focus specifically on CAR-associated signaling cascades. To assess whether signaling influenced the establishment of infection, we measured levels of HIV-1 proviral DNA using an established nested *Alu*-PCR approach [20]. We modified the assay by designing primers to luciferase to estimate the relative frequency of HIV integration without confounding signals from the lentiviral vectors used to express the CARs (see materials and methods). CAR-associated signaling did not affect the infection of Jurkat cells since we detected comparable levels of provirus regardless of the presence or absence of CAR ligand (Fig 2A). When HIV-1 expression was measured by luciferase activity, Jurkat cells infected in the context of strong T cell signaling expressed greater than 10-fold more HIV-1 compared to untreated controls (Fig 2B). In contrast, engagement of the low affinity receptor led to a modest 3-fold expression compared to unstimulated cells despite a similar proviral load as the high affinity CAR-expressing cells. These data indicate that strong T cell signaling at the time of infection facilitates HIV-1 expression without enhancing provirus integration.

**Fig 2.**
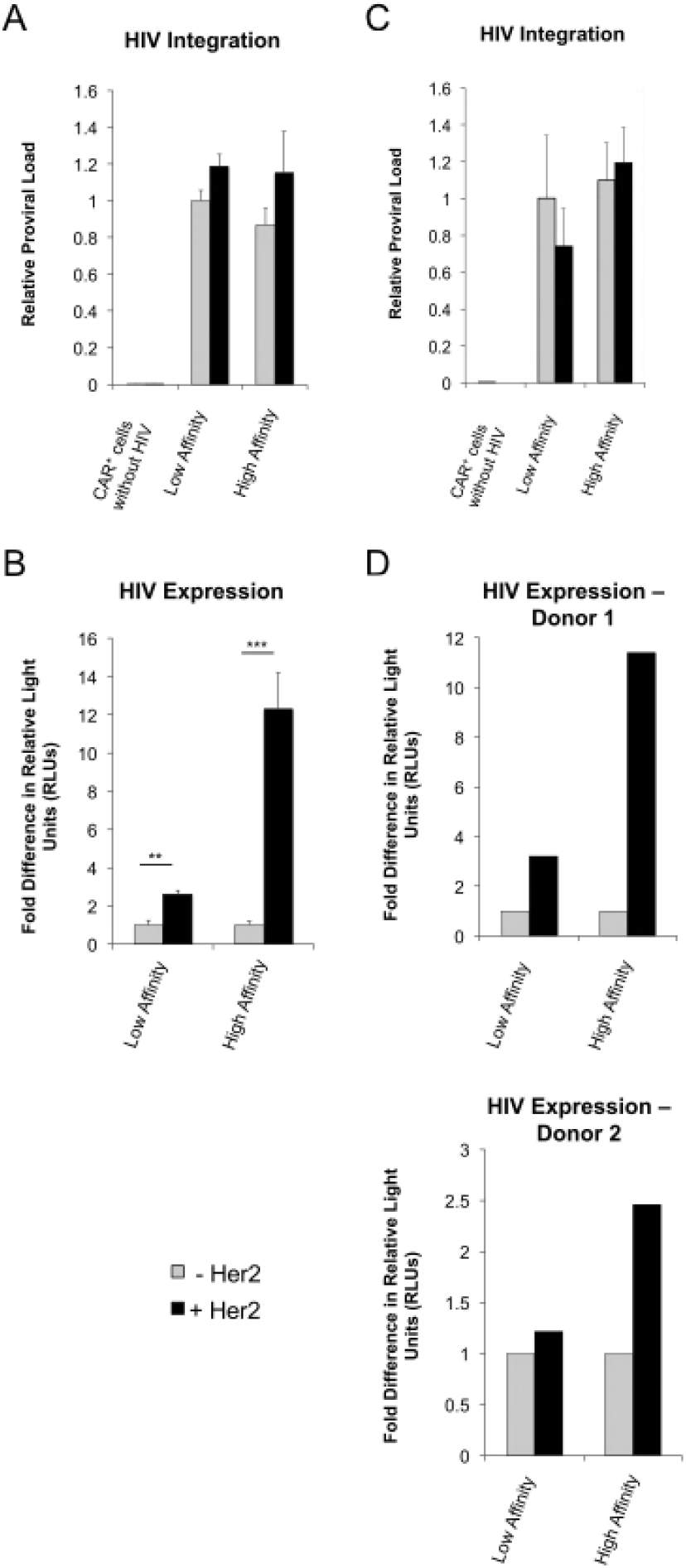
T cell signaling at the time of HIV-1 infection regulates provirus expression. (*A* and *B*) Jurkat T cells were transduced with the high or low affinity CAR. Cells were stimulated through the CAR at the time of HIV-1 infection with VSV-G pseudotyped NL4–3.Luc. (*A*) Relative levels of integrated provirus 24 h post infection of high or low affinity Jurkat T cells using nested *Alu*-PCR. Uninfected CAR-expressing Jurkat T cells were a negative control. (*B*) Luciferase activity measured 24 h post-infection presented as fold difference in relative light units (RLUs) over unstimulated cells for each CAR+ cell line. **p<0.005,***p≤0.0005. *A* and *B* were performed in triplicate and are representative of four independent experiments. Data are presented as mean ± standard deviation. (*C* and *D*) Primary CD4^+^ T cells isolated from healthy human donors were transduced with CARs and given one week to return to a resting state. Cells were stimulated through the CAR at the time of HIV infection with single-round VSV-G pseudotyped NL4–3.Luc. (*C*) Relative levels of integrated provirus 24 h after infection of high or low affinity CAR-expressing primary T cells (from the same donor) using nested *Alu*-PCR. Uninfected CAR-expressing primary T cells were a negative control. Results are from a single experiment performed in triplicate and are representative of two independent experiments. Data are presented as mean ± standard deviation. (*D*) Luciferase activity measure 4 days post-infection presented as fold difference in RLUs over unstimulated cells for each CAR+ cell line. Experiments from two separate donors are shown and are representative of three independent experiments. See also S2 Fig.

We confirmed that these differences were due to downstream signaling emanating from the CARs by using the src kinase inhibitor PP2. In the presence of PP2, the increase in HIV-1 expression upon cellular stimulation was attenuated, consistent with T cell signaling as a regulator of HIV-1 expression (S2 Fig). The pharmacologically inactive version of this inhibitor, PP3, had no effect on the ability of CARs to influence HIV-1 expression.

We validated these results using primary CD4+ T cells that were transduced with either the low affinity or high affinity CAR. Following transduction, cells were allowed to return to a resting state as monitored by low CD69 expression before infection with HIV-1 in the absence or presence of the ligand Her2. Consistent with the data from Jurkat cells, similar levels of proviral DNA were detected in primary T cells regardless of CAR signaling (Fig 2C). Cells that received robust stimulation at the time of infection expressed 1.5- to 4-fold more HIV-1 than cells stimulated through the low affinity receptor (Fig 2D).

To gain insight into whether there is a threshold or minimal T cell signal required for HIV-1 infection and replication, we transduced Jurkat T cells with CAR receptors that spanned a range of binding affinities (Fig 1B). These cells were infected with NL4–3.Luc as described above in the absence or presence of Her2. Although the high affinity condition supported HIV-1 infection and transcription, the intermediate and low affinity receptors did not support HIV-1 expression (Fig 3A). This was despite similar levels of infection as determined by measuring proviral DNA (Fig 3B). These data suggest that T cell signaling controls HIV-1 expression by a digital on/off mechanism since viral expression does not linearly correlate with signal strength.

**Fig 3.**
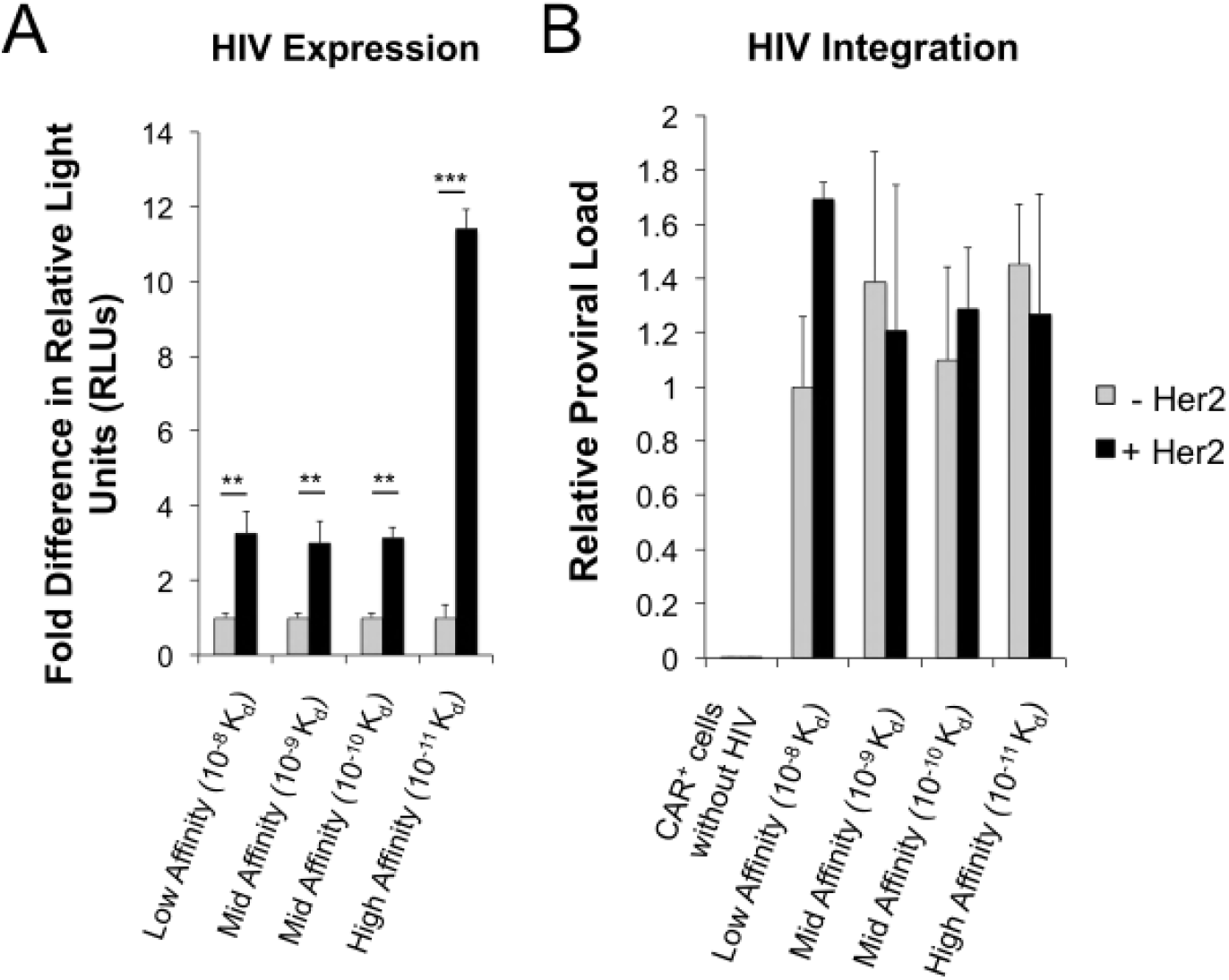
Robust T cell signaling is required for HIV transcription. (*A*) Jurkat T cells were transduced and positively selected for indicated CARs and then infected with NL4–3.Luc. 24 h post infection, cells were lysed for luciferase analysis. Data are presented as fold difference in RLUs over unstimulated cells for each CAR+ population. **p<0.005, ***p<0.0001. *B*) Relative levels of integrated provirus after infection of CAR-expressing Jurkats using nested *Alu*-PCR. *A* and *B* were performed in triplicate and are representative of four independent experiments. Data are presented as mean ± standard deviation.

### Robust signals during HIV-1 infection establishes an inducible latent reservoir

We hypothesized that differential T cell signaling during infection alters the size of the inducible latent reservoir. To examine this, we infected CAR-expressing primary CD4+ T cells with VSV-G pseudotyped BRU-dENV-GFP in the presence of Her2 ligand. One week post infection, cells were sorted for both mCherry expression as a marker for the CAR and lack of GFP expression in order to enrich for latently infected cells. CAR^+^/GFP^neg^ cells were reactivated with PMA plus ionomycin or left unstimulated to control for spontaneous HIV-1 reactivation (Fig 4A). PMA plus ionomycin significantly reactivated HIV-1 expression within cells that had been initially infected in the context of strong signaling, resulting in a 3- to 9-fold increase in the percentage of GFP positive cells and a 1000-fold induction of HIV-1 mRNA measured by qRT-PCR (Fig 4B). However, the observed reactivation of HIV-1 was modest in cells infected at the time of stimulation through the low affinity CAR. A 200-fold induction of HIV-1 RNA was detected in reactivated latently infected cells expressing low affinity CARs, and less than a 2-fold change was observed in the percentage of GFP^+^ cells. Therefore, despite both minimal and robust signaling resulting in comparable amounts of integrated HIV-1 provirus, robust signaling was not only necessary for active transcription but also supported the generation of a population of latently infected cells that could be readily induced to express HIV-1. The population of latent cells generated in response to weaker CAR signaling was more resistant to reversal suggesting that HIV-1 in these cells was strongly repressed.

**Fig 4.**
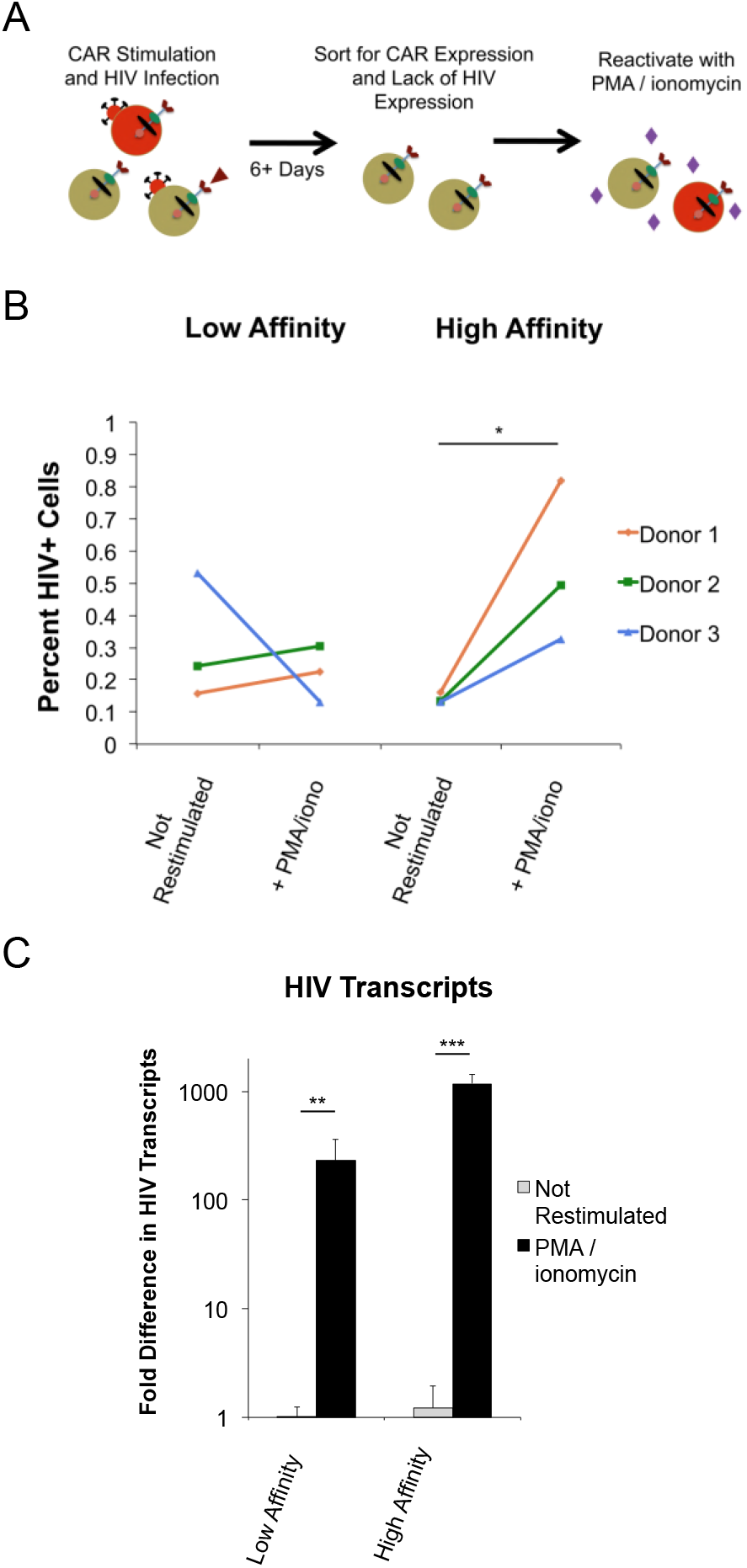
Robust signals during HIV-1 infection establishes an inducible latent reservoir. (*A*) Outline of experimental plan to enrich for latently infected cells following infection. Primary CD4+ T cells are infected with BRU-deltaEnv-GFP and GFP-negative cells are sorted to enrich for latently infected cells. (*B*) Percent GFP+ HIV-expressing cells after stimulation of latent cells with PMA plus ionomycin or without stimulation. Data are from three separate donors. (*C*) Latently-infected cells were restimulated with PMA and ionomycin. HIV-1 expression was monitored by measuring Tat RNA by qRT-PCR. Values are shown as fold difference in HIV transcripts over corresponding non-reactivated controls. Note the log scale. All data in *C* are derived from 4–6 replicates and are representative of three independent experiments with different donors. Data are presented as mean ± standard deviation. *p<0.05, **p<0.005, ***p<0.0001.

### RNAP II Processivity Limits HIV-1 Transcription in the Absence of Robust Signaling

We were interested in mechanisms that governed HIV-1 repression following integration in the absence of sufficient T cell signaling; therefore, we examined the binding of transcriptional regulators on the HIV-1 LTR by chromatin immunoprecipitation (ChIP). Jurkat T cells expressing low or high affinity CARs were infected with NL4-3.Luc in the absence or presence of Her2 ligand. One day post-infection, cells were fixed and chromatin was prepared for ChIP.

Since HIV-1 proviral latency correlates with a positioned nucleosome that is downstream of the transcriptional start site, we explored whether the LTR was associated with post-translationally modified histones as an indicator of chromatin organization. ChIPs for acetylated histone H3 showed no significant difference in binding of the HIV-1 LTR between cells infected in the absence or presence of T cell signaling (Fig 5A). Therefore, chromatin accessibility does not appear to be limiting HIV-1 proviral transcription following infection.

**Fig 5.**
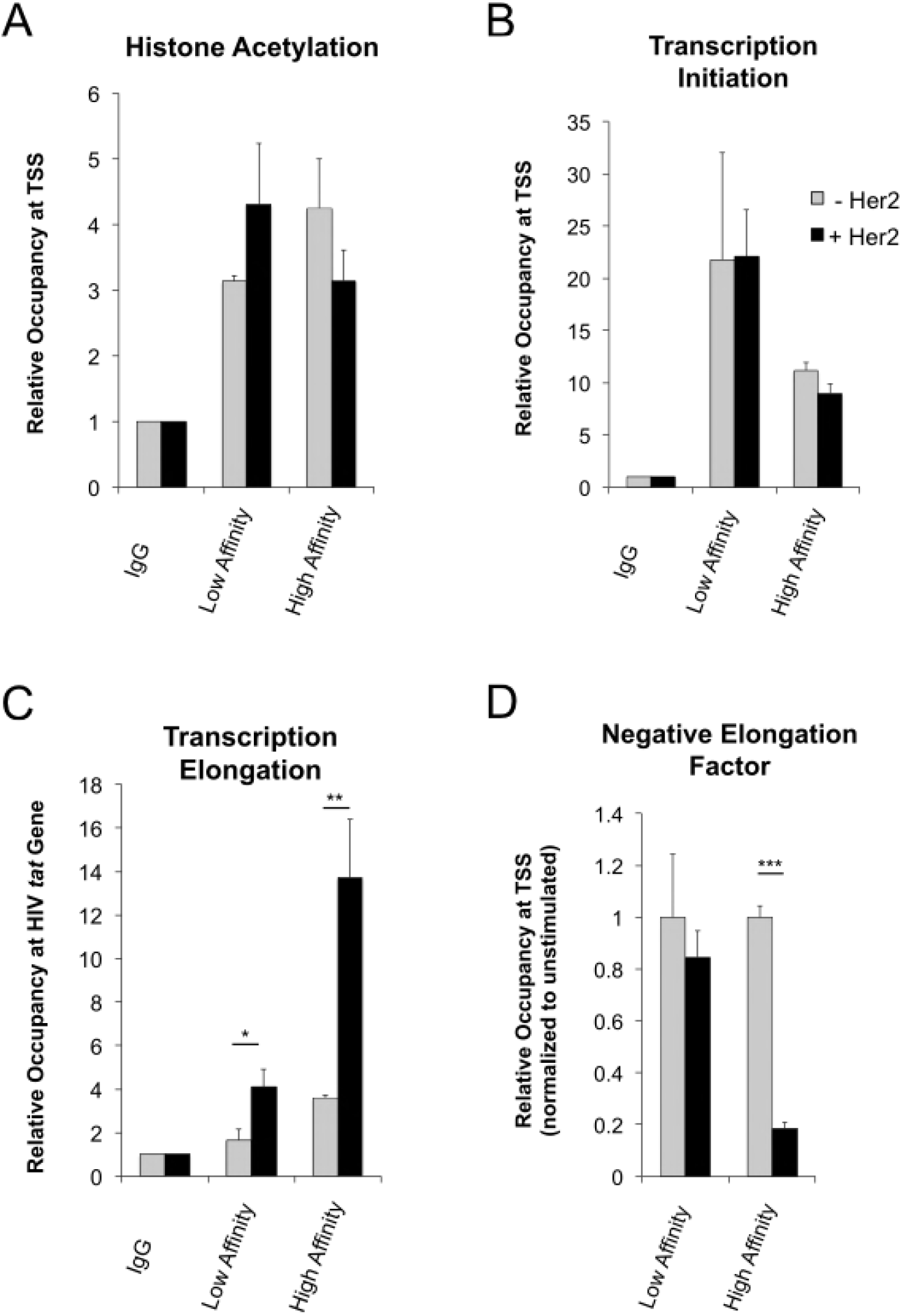
RNAP II Processivity Limits HIV-1 Transcription in the Absence of Robust Signaling. (*A*) ChIP for presence of acetylated H3 near the transcriptional start site (nuc1). (*B*) ChIP for RNAP II at the HIV transcriptional start site. (*C*) ChIP for RNAP II associated with the HIV *tat* gene to measure polymerase processivity. Data from *A*-*C* are normalized to corresponding IgG controls for each stimulation condition. (*D*) ChIP for NELF-d at the HIV transcriptional start site. Data is normalized to corresponding unstimulated condition for each CAR+ cell line. *A*-*D* were performed in triplicate and are representative of at least three independent experiments in Jurkat T cells. Data are presented as mean ± standard deviation. Primers used for HIV transcriptional start site are +30 and +239. Primers used for *tat* gene are +5379 and +5482. *p<0.05, **p<0.005, ***p≤0.0005.

We then examined RNAP II processivity by measuring RNAP II occupancy at multiple points, including the transcriptional start site and downstream in the HIV *tat* gene. RNAP II was detected at the HIV transcriptional start site whether cells were activated through a CAR or were unstimulated (Fig 5B). However, signaling through the high affinity receptor resulted in an increase in downstream RNAP II by greater than 4-fold, whereas only modest levels of RNAP II were found downstream in the absence of signals or following weak signaling (Fig 5C).

Since these data indicated a role for transcriptional pausing, we examined if CAR signaling altered the presence of the pausing factor NELF at the HIV-1 transcriptional start site. Using ChIPs, we determined that signaling through the high affinity receptor diminished binding of NELF at the HIV LTR by greater than 85% (Fig 5D). These data support a model in which a lack of robust T cell signaling limits HIV transcription by establishing a paused polymerase complex.

## Discussion

Previous studies suggest that cell signaling may be a key regulator of HIV-1 expression and latency. The latent reservoir is enriched for antigen specific T cells, including those that respond to CMV, HSV, tuberculosis, and HIV [21–25]. Furthermore, the use of superantigens during viral entry increases HIV-1 replication [26]. Partial activation, cellular polarization, cell-to-cell contact, and/or infection of resting quiescent cells through perturbation have also been suggested to bias infections towards latency [11,27–31]. Therefore, the extent of cell activation is a key determinant in regulating the course of HIV-1 infection including the formation of the reservoir.

We have shown that differential signaling through CARs, which mimic TCR signaling, influences HIV-1 transcription and latency. In the lymph node, a primary site for both HIV replication and the persistent latent reservoir [32–34], T cells will sample lymph node resident cells in search for antigen. Some of these interactions, facilitated by the presentation of the T cell cognate antigen, will result in robust T cell activation, clonal expansion, and changes in gene expression. However, most MHC complexes will lack cognate antigen and initiate weak signaling [35,36]. Using multiple CARs whose affinities for the Her2 ligand span several logs, we can deliver a range of signaling inputs to model the spectrum of T cell receptor signaling events. Our data indicates that stronger T cell activation at the time of infection, which would be more similar to antigen specific responses, correlates with robust HIV-1 expression as well as the establishment of inducible latently infected cells.

Having a library of CARs with a range of binding affinities allowed us to determine if HIV-1 responds to signaling in an analog fashion correlating with signal input or digitally regulated by specific thresholds resulting in all-or-none responses [37]. Signaling through the CARs with affinities that were intermediate did not support active transcription despite a greater than 10-fold increase in binding affinity compared to our low affinity receptor. These results would suggest that TCR signaling provides more of an on/off switch and that there exist signaling thresholds that must be overcome to assure efficient transcription and replication.

Signal transduction and gene expression are inherently noisy processes, and stochastic events are hypothesized to drive HIV latency. That latency and HIV replication are driven by episodic bursts of proviral transcription and Tat levels has been supported by mathematical modeling and experiments using engineered virus models [38–40]. Even if latency is driven by random fluctuations of provirus transcription, T cell associated signals are strong modulators of noise, and targeting these pathways could enhance treatments directed at HIV reactivation [41]. However, it is important to appreciate that although signaling and transcription are subject to stochastic variation, these are coordinated and combinatorial processes that lead to defined patterns of gene expression and phenotypic outcomes [42].

Regulated aspects of transcription include assembly of multi-subunit complexes such as RNAP II and associated cofactors, chromatin, and transcription factors at the LTR. Our data suggest that the association of NELF with RNAP II is regulated by TCR signaling. Multiple positive and negative signals are known to converge on NELF-driven transcriptional pausing. P-TEFb relieves NELF repression through phosphorylation [43] and is itself regulated by cellular stress and signals [44–46]. Furthermore, we have shown that NELF interacts with co-repressors including NCoR1-GPS2-HDAC3 at the HIV-1 promoter [47] which may reinforce HIV latency, especially during chronic infection, by facilitating post-translational modifications of histones and chromatin organization.

We propose that strength of signal at the time of infection acts as a bifurcating event leading to either robust transcription and the establishment of an inducible latent reservoir *or* minimal transcription and deep-seated latency. Our observations are consistent with the previous characterization of patient reservoirs that identified three subsets of latently infected cells: a small population of cells carrying inducible provirus, a larger populations of cells with intact proviruses that are difficult to reactivate, and many defective proviruses [48]. Successful purging of the latent reservoir may require the use of a cocktail of latency reversing agents or the development of novel strategies to block reactivation [49–51].

## Materials and Methods

### Cells

Jurkat CD4+ T cells (E6–1) and human embryonic kidney 293T cells were obtained from American Type Culture Collection (ATCC). Jurkat cells were cultured in RPMI 1640, 5% FBS (Corning, Inc.), 100^units^/_mL_ penicillin (Invitrogen), 100^μ^g/_mL_ streptomycin (Invitrogen), and 2mM L-glutamine (Invitrogen). 293T cells were cultured in Dulbecco’s Modified Eagle Medium, 10% FBS, 100^units^/_mL_ penicillin, 100^μ^g/_mL_ streptomycin, and 2mM L-glutamine. Cells were grown at 37° C with 5% CO_2_.

Primary CD4+ T cells were derived from de-identified healthy blood leukapheresis packs purchased from NY Biologic. Mononuclear cells were enriched from leukapacks by centrifugating through Histopaque gradient (Sigma-Aldrich). CD4+ T cells were isolated by negative selection using EasySep Human CD4+ T Cell Enrichment Kits from StemCell Technologies. CD4+ cells were maintained in RPMI 1640, 10% FBS, 100^units^/_mL_ penicillin, 100^μg^/_mL_ streptomycin, and 2mM L-glutamine at 37° C with 5% CO_2_. Prior to transduction with CARs, primary cells were supplemented with 10^units^/_mL_ IL-2 and 10^ng^/_mL_ IL-7. Following transduction, IL-2 was removed from culture conditions. All cells and cell lines were split every 2–3 days.

### Viruses and transductions

CARs were driven by a SFFV promoter in the lentiviral vector pHR [18,19]. pNL4–3.Luc.R-E-was obtained from NIH AIDS Reagent Program. BRU-ΔEnv-GFP was a gift from Gregory Viglianti (Boston University). Lentiviruses were made by transfection of vectors, VSV-G, Rev, Tat, and Gag-Pol constructs into 293T cells with 45μL polyethylenimine (1^mg^/_mL_) per 6×10^6^ cells. Supernatants were collected, filtered with 0.45μm syringe filter (Corning), concentrated by centrifuging through a 20% sucrose gradient, and titered with CEM cells [52]. We used a range of multiplicity of infections, but most viruses and lentiviruses within this paper were concentrated to approximately 1×10^6IU^/_mL_.

For transductions with CAR vectors, primary and Jurkat cells were stimulated for 5–6 h with 10 ^μg^/_mL_ PHA, washed in PBS, and spinoculated with lentivirus and 5^μg^/_mL_ polybrene (Millipore) at 1200g for 90 min. Cells were then supplemented with fresh RPMI and IL-7, cultured overnight, and washed in PBS 18 h later. Cells were rested for one week to return to a resting state as confirmed by low CD69 expression (Brilliant Violet 421 anti-human CD69 antibody; Clone FN50, BioLegend) prior to HIV-1 infection.

### CAR stimulation and infections

Non-tissue culture treated plates were coated overnight at 37°C with 1^μ^g/_mL_ Her2 (Recombinant Human ErbB2/Her2 Fc Chimera Protein from R&D Systems, 1129-ER). Her2 solution was removed from wells, plates were washed 3 times in PBS, and wells were blocked for 1 h with a 5% FBS-PBS solution.

Jurkat or primary CD4+ T cells were infected and simultaneously plated in Her2-treated wells. For experiments in which latently infected cells were generated, cells were spinoculated in the Her2-treated wells at 1200xg for 90 min and then supplemented with fresh RPMI and IL-7. Following overnight infection, cells were washed and either lysed or maintained in fresh media in the absence of Her2.

For reactivation of latent cells, mCherry (CAR) positive and GFP (HIV) negative cells were sorted at 6 or 7 days post HIV infection. Cells were cultured with 5^ng^/_mL_ PMA and either 10 or 100uM ionomycin for 2.5 h. Reactivated cells were washed in PBS and re-plated in media supplemented with 10^ng^/_mL_ IL-7. Cells were cultured overnight prior to fixation for flow analysis. For some experiments, cells were treated with 10 μM PP2 or PP3 (Calbiochem – Millipore Sigma) at the time of infection.

### Luciferase analysis

Jurkat cells were washed and lysed for luciferase analysis 24 h post infection, while primary T cells were measured at 4 days post infection. Luciferin (Promega) was added and luciferase activity was measured via BioTek Synergy HT Microplate Reader.

### Nested *Alu*-PCR

Nested PCR strategy was adapted from Agosto *et al*., 2007 [20]. Briefly, integrated HIV DNA was amplified using forward primers for the luciferase sequence and reverse primers for human *Alu* (see Table 1). The first reaction was performed on a TProfessional Thermocycler from Biometra according to the following conditions: 4 m at 95° followed by 20 cycles of 15 s at 93°, 15 s at 50°, and 2.5 m at 70°. A second round of amplification was then performed using a forward primer, a reverse primer, and a probe for real time PCR within the HIV-1 3’ R / U5 region (see Table 1). The amount of amplified copies of HIV was determined based on an NL4-3 plasmid copy standard. The second reaction was performed on an Applied Biosystems QuantStudio 3 Real-Time PCR system with heating for 4 m at 95° and real-time PCR conditions of denaturation for 15 s at 95°, annealing for 30 s at 60°, and extension for 1 m at 72°.

**Table 1.**
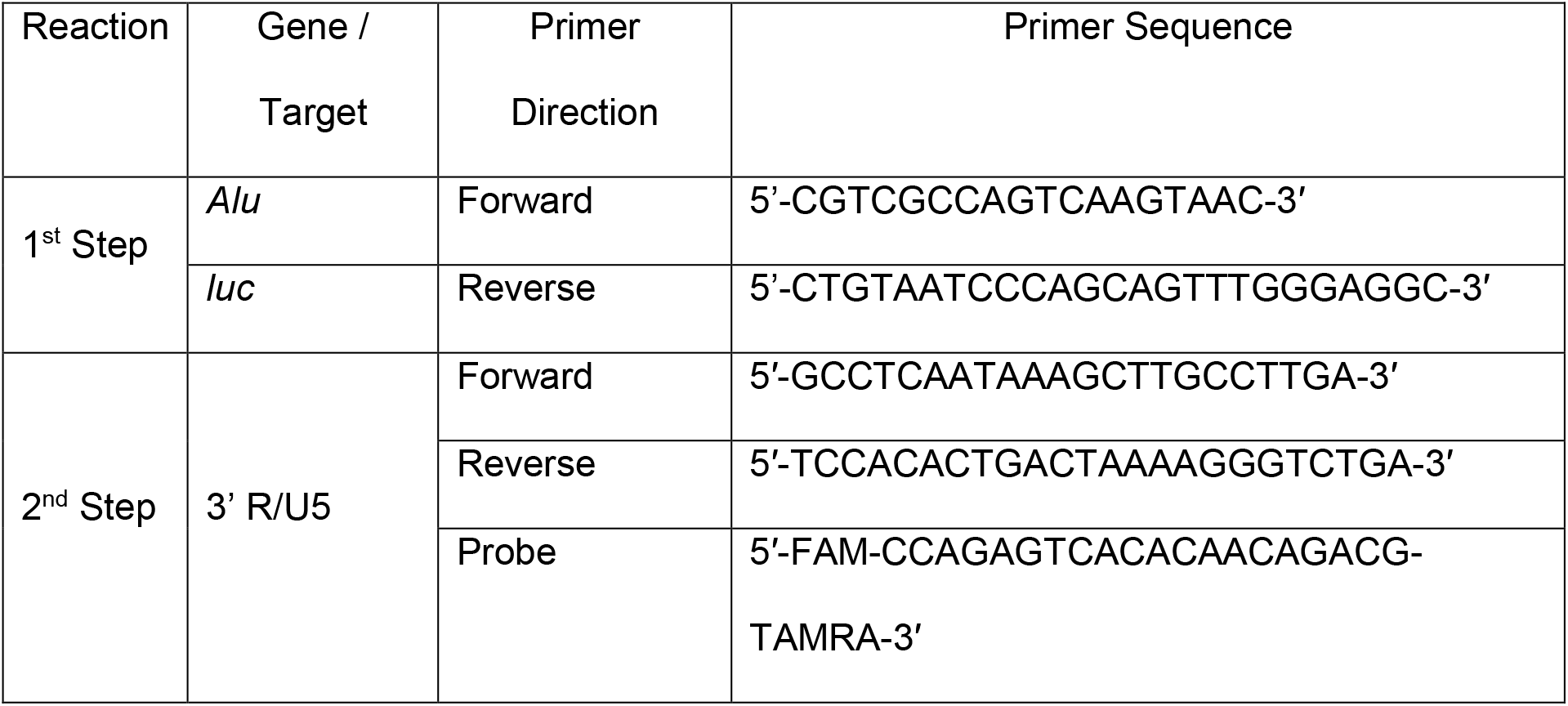
Oligos used for *Alu*-PCR.

### Flow cytometry

Flow data were collected on an LSRII from BD Biosciences. Zombie UV Fixable Viability Kit (BioLegend) was used as live/dead stain for reactivation experiments. All cells were washed and fixed in a final concentration of 2% paraformaldehyde prior to analysis. Cell sorting was performed on a MoFlo Astrios from Beckman Coulter. All flow experiments performed at Boston University School of Medicine Flow Cytometry Core Facility.

### RT-PCR

RT-PCR for HIV-1 mRNA was performed using forward primers and reverse primers for unspliced HIV Tat sequence, and all values were normalized against beta-actin as a housekeeping gene (see Table 2). The second reaction was performed on an Applied Biosystems QuantStudio 3 Real-Time PCR system with heating for 15 m at 94° and real-time PCR conditions of denaturation for 15 s at 94°, annealing for 30 s at 60°, and extension for 30 s at 72°.

**Table 2.**
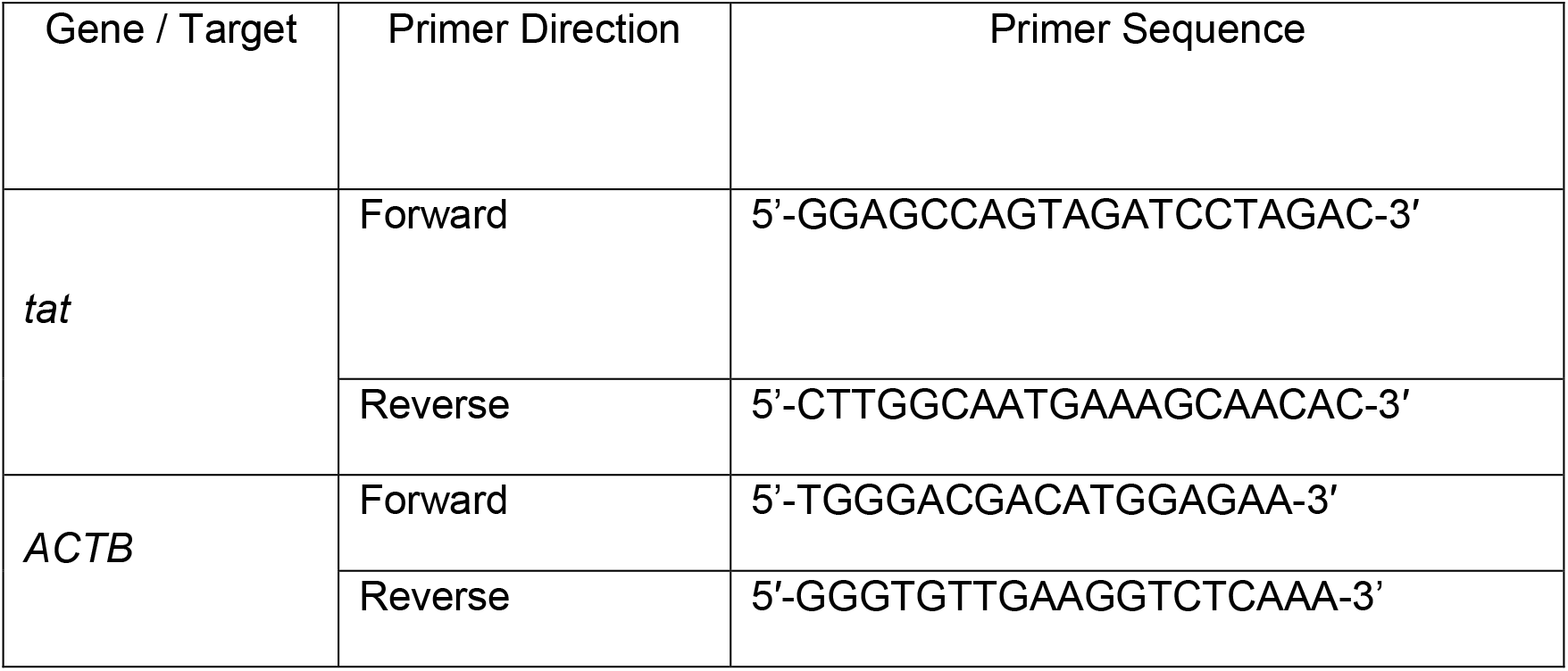
Oligos used for RT-PCR.

### Chromatin immunoprecipitation

ChIP was performed according to Natarajan *et al*., 2013 [47] with the addition of a nuclei isolation step using Farnham Lysis Buffer prior to sonication. Briefly, cells were washed in PBS and fixed in a final concentration of 1% formaldehyde in methanol. Crosslinking was quenched by addition of glycine to a final concentration of 240mM. Cells were then washed, centrifuged, and flash frozen in liquid nitrogen. Pellet was lysed in Farnham Lysis Buffer (5mM PIPES pH 8.0, 85mM KCl, 0.5% NP-40 with addition of Halt Protease and Phosphatase Inhibitor Single-Use Cocktail from ThermoFisher) before centrifugation to obtain nuclei fraction. Nuclei were lysed in RIPA buffer prior to sonication in BioRupter Pico for 15 m with alternating 30 s cycles. Samples were centrifuged to remove debris and pre-cleared with addition of 50% protein A sepharose bead for 30 m at 4°C. Beads were then pelleted and supernatants were split into 100μL portions as input DNA and multiple 300μL portions for sample analysis. Samples were incubated with antibodies overnight at 4°C. Antibody-bound proteins and cross-linked DNA were isolated by addition of 50% protein A sepharose beads for 2 h at 4°C prior to centrifugation.

Immunoprecipitates were then washed with low salt (0.1% SDS, 1% Triton X-100, 2mM EDTA, 20mM Tris-HCl pH 8.0, 150mM NaCl), high salt (0.1% SDS, 1% Triton X-100, 2mM EDTA, 20mM Tris-HCl pH 8.0, 500mM NaCl), lithium wash (0.25M LiCl, 1% NP-40, 1% sodium deoxycholate, 1mM EDTA, 10mM Tris-HCl), and Tris-EDTA buffers (10mM Tris, 1mM EDTA) before use of elution buffer (1% SDS, 0.1M NaHCO_3_). Cross-linking was reversed with addition of 5M NaCl overnight at 65°C to both samples and input DNA before addition of proteinase K to isolate DNA. Sample DNA was purified using ChIP DNA Clean & Concentrator Kit (Zymo Research).

Antibodies used included anti-NELF-d (Antibody TH1L from Proteintech Group), anti-RNA Polymerase II antibody (Clone N20 from Santa Cruz Biotechnology), anti-histone H3 antibody (Product 06–599 from Millipore Sigma), and Normal Rabbit IgG (Product 12–370 from Millipore Sigma).

Primers used for the transcriptional start site include the forward primer at +30 and the reverse primer at +239. Primers used for transcriptional elongation include the forward and reverse primers within the *tat* gene (see Table 3).

**Table 3.**
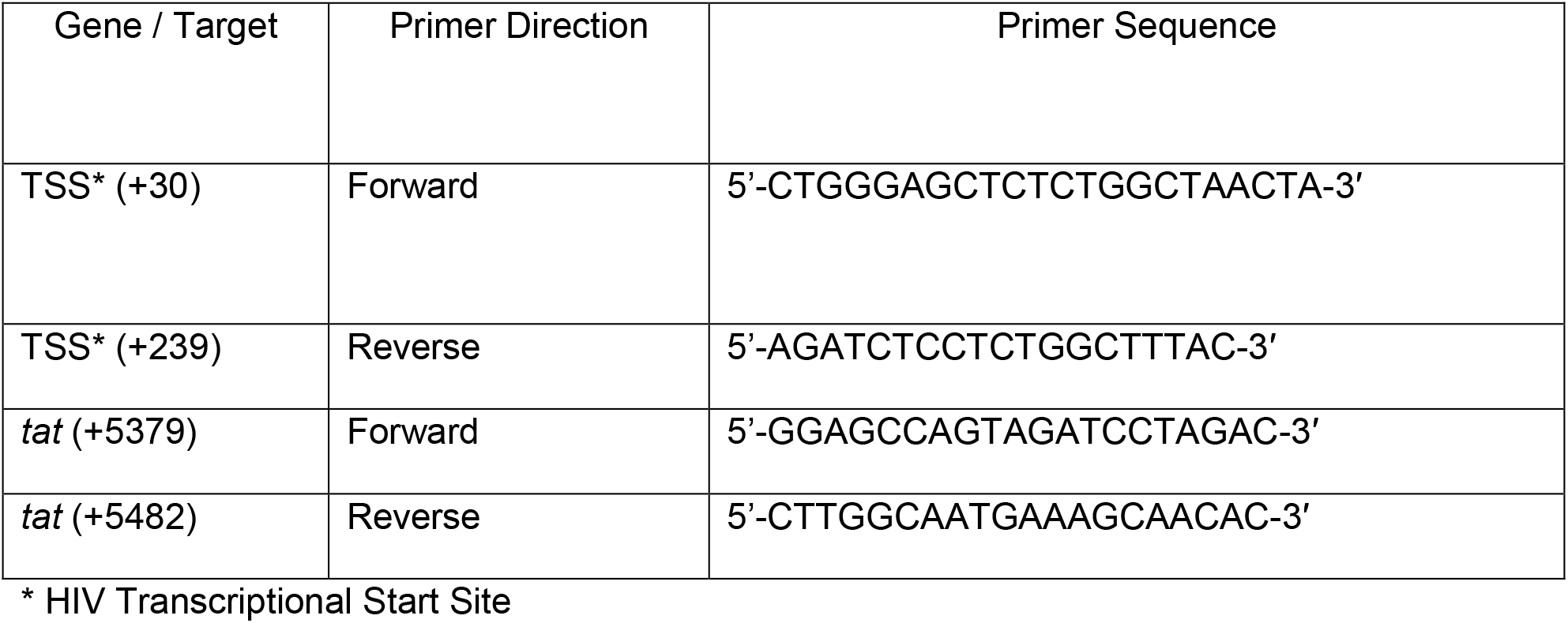
Oligos used for Chromatin Immunoprecipitation Assays.

### Statistical analysis

All statistical analysis performed using unpaired student’s *t* test.

## Acknowledgements

We would like to thank the Boston University School of Medicine Flow Cytometry Core Facility for their contributions to the flow analysis and cell sorting experiments. Melissa Herring assisted with the kinase inhibitor assays. Kyle Pedro, Luis Agosto, PhD, and Gregory Viglianti, PhD (Boston University School of Medicine) all provided advice regarding experimental design and manuscript publication.

## Supporting information legends

**S1 Fig.**
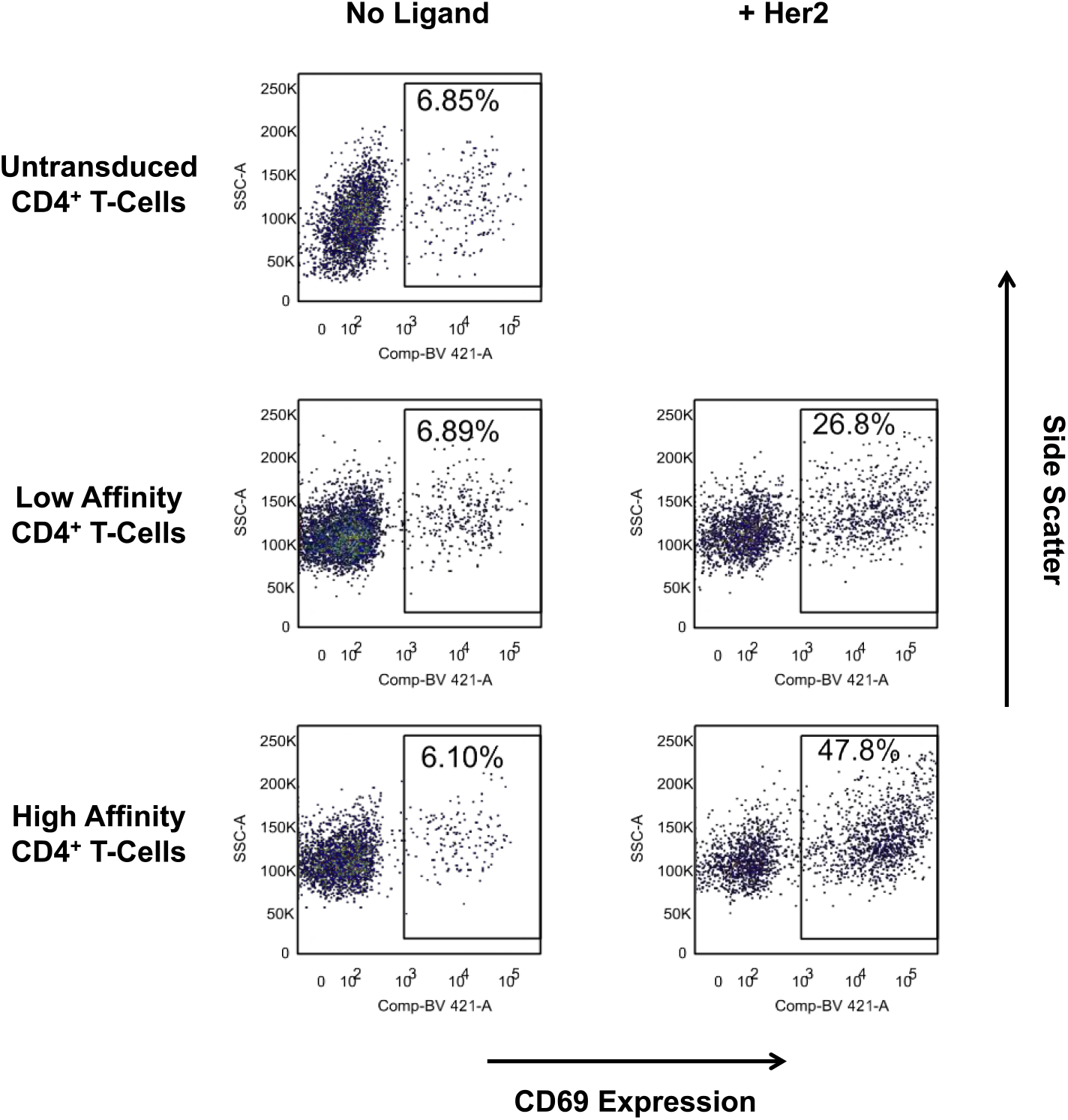
Differential signaling through chimeric antigen receptors. Related to Fig 1. CD4+ T cells isolated from healthy human donors were transduced with low affinity or high affinity CARs and then allowed to return to a resting state. Cells were either then stimulated through the receptor or left unstimulated. CD69 expression was compared to a negative control of untransduced cells one day after Her2 stimulation. Data is presented as dot plots based on flow cytometry analysis and is from a representative experiment that has been performed four times with different donors.

**S2 Fig.**
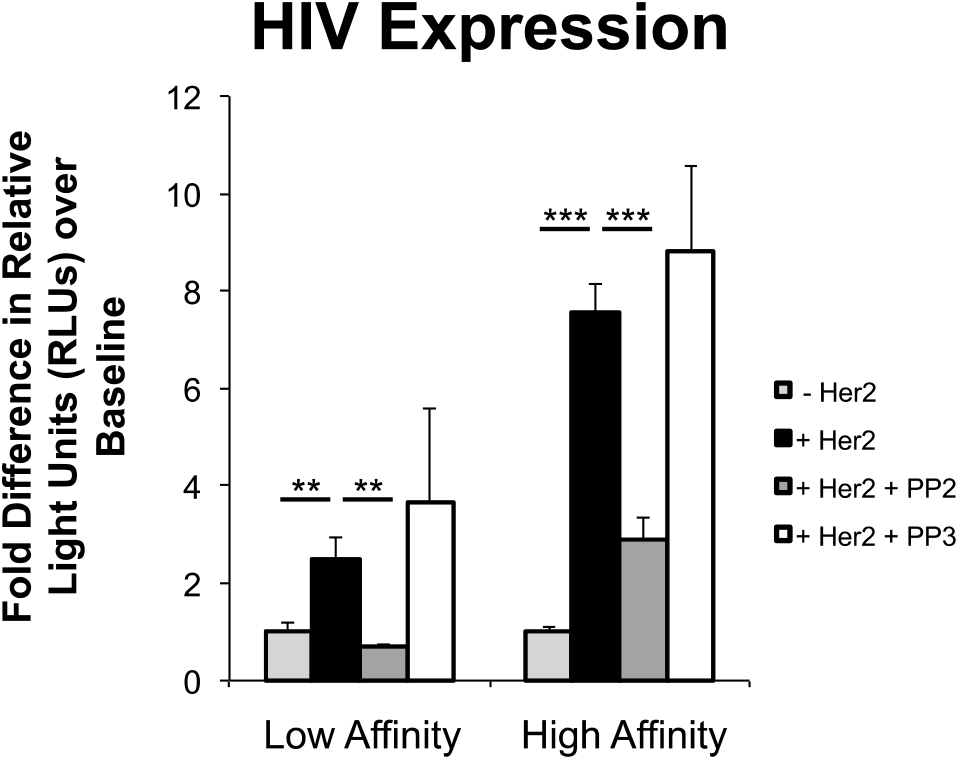
Src kinase inhibitor PP2 inhibits CAR-mediated HIV transcription. Related to Fig 2. CAR+ Jurkat T cells were stimulated with or without Her2 in the absence or presence of 10 μM PP2 or PP3 at the time of HIV infection with single-round VSV-G pseudotyped NL4–3.Luc. 24 h post infection, cells were lysed to measure luciferase. Data are presented as fold difference in RLUs over unstimulated cells for each CAR+ population. **p<0.005, ***p≤0.0005. S2 Fig was perfomed in triplicate and is representative of five independent experiments. Data are presented as mean ± standard deviation.

